# A subset of pediatric thalamic gliomas share a distinct DNA methylation profile, H3K27me3 loss and frequent alteration of *EGFR*

**DOI:** 10.1101/2020.08.28.239160

**Authors:** Philipp Sievers, Martin Sill, Daniel Schrimpf, Damian Stichel, David E. Reuss, Dominik Sturm, Jürgen Hench, Stephan Frank, Lenka Krskova, Ales Vicha, Michal Zapotocky, Brigitte Bison, Patrick N. Harter, Matija Snuderl, Christof M. Kramm, Guido Reifenberger, Andrey Korshunov, Nada Jabado, Pieter Wesseling, Wolfgang Wick, David A. Solomon, Arie Perry, Thomas S. Jacques, Chris Jones, Olaf Witt, Stefan M. Pfister, Andreas von Deimling, David T. W. Jones, Felix Sahm

**Affiliations:** Department of Neuropathology, Institute of Pathology, University Hospital Heidelberg, Heidelberg, Germany; Clinical Cooperation Unit Neuropathology, German Consortium for Translational Cancer Research (DKTK), German Cancer Research Center (DKFZ), Heidelberg, Germany; Hopp Children’s Cancer Center Heidelberg (KiTZ), Heidelberg, Germany; Division of Pediatric Neurooncology, German Cancer Consortium (DKTK), German Cancer Research Center (DKFZ), Heidelberg, Germany; Pediatric Glioma Research Group, German Cancer Research Center (DKFZ), Heidelberg, Germany; Department of Pediatric Oncology, Hematology, Immunology and Pulmonology, University Hospital Heidelberg, Heidelberg, Germany; Institute for Medical Genetics and Pathology, University Hospital Basel, Basel, Switzerland; Prague Brain Tumor Research Group, Second Faculty of Medicine, Charles University and University Hospital Motol, Prague, Czech Republic; Department of Pathology and Molecular Medicine, Second Faculty of Medicine, Charles University and University Hospital Motol, Prague, Czech Republic; Department of Pediatric Haematology and Oncology, Second Faculty of Medicine, Charles University and University Hospital Motol, Prague, Czech Republic; Institute of Diagnostic and Interventional Neuroradiology, University Hospital Würzburg, Würzburg, Germany; Institute of Neurology (Edinger Institute), Goethe University, Frankfurt, Germany; German Cancer Consortium (DKTK), Partner Site Frankfurt/Mainz, Frankfurt am Main, Germany; German Cancer Research Center (DKFZ), Heidelberg, Germany; Frankfurt Cancer Institute (FCI), Frankfurt am Main, Germany; Department of Pathology, NYU Langone Medical Center, New York, NY, USA; Division of Pediatric Hematology and Oncology, University Medical Center Göttingen, Göttingen, Germany; Institute of Neuropathology, Heinrich Heine University, Düsseldorf, Germany; German Cancer Consortium (DKTK), Partner Site Essen/Düsseldorf, Germany; Department of Human Genetics, McGill University, Montreal, QC H3A 1B1, Canada; Department of Pediatrics, McGill University, Montreal, QC H4A 3J1, Canada; The Research Institute of the McGill University Health Center, Montreal, QC H4A 3J1, Canada; Department of Pathology, Amsterdam University Medical Centers, Location VUmc and Brain Tumor Center Amsterdam, Amsterdam, The Netherlands; Princess Máxima Center for Pediatric Oncology, Utrecht, The Netherlands; Clinical Cooperation Unit Neurooncology, German Consortium for Translational Cancer Research (DKTK), German Cancer Research Center (DKFZ), Heidelberg, Germany; Department of Neurology and Neurooncology Program, National Center for Tumor Diseases, Heidelberg University Hospital, Heidelberg, Germany; Department of Pathology, University of California, San Francisco, CA, USA; Clinical Cancer Genomics Laboratory, University of California, San Francisco, CA, USA; Department of Neurological Surgery, University of California, San Francisco, CA, USA; Developmental Biology and Cancer Research and Teaching Department, UCL Great Ormond Street Institute of Child Health, London, UK; Department of Histopathology, Great Ormond Street Hospital for Children NHS Foundation Trust, London, United Kingdom; Division of Molecular Pathology, Institute of Cancer Research, London, United Kingdom; Clinical Cooperation Unit Pediatric Oncology, German Cancer Research Center (DKFZ) and German Consortium for Translational Cancer Research (DKTK), Heidelberg, Germany

**Keywords:** Pediatric high-grade glioma, (bi-)thalamic, DNA methylation profile, *EGFR* mutation, H3 K27M mutation, K27me3

## Abstract

**Background:** Malignant astrocytic gliomas in children show a remarkable biological and clinical diversity. Small in-frame insertions or missense mutations in the *EGFR* gene have recently been identified in a distinct subset of pediatric bithalamic gliomas with a unique DNA methylation pattern.

**Methods:** Here, we investigated an epigenetically homogeneous cohort of malignant gliomas (n=58) distinct from other subtypes and enriched for pediatric cases and thalamic location, in order to elucidate the overlap with this recently identified subtype of pediatric bithalamic gliomas.

**Results:** *EGFR* gene amplification was detected in 16/58 (27%) tumors, and missense mutations or small in-frame insertions in *EGFR* were found in 20/30 tumors with available sequencing data (67%; five of them co-occurring with *EGFR* amplification). Additionally, eight of the 30 tumors (27%) harbored an H3.1 or H3.3 K27M mutation (six of them with a concomitant *EGFR* alteration). All tumors tested showed loss of H3K27me3 staining, with evidence of *EZHIP* overexpression in the H3 wildtype cases. Although some tumors indeed showed a bithalamic growth pattern, a significant proportion of tumors occurred in the unilateral thalamus or in other (predominantly midline) locations.

**Conclusions:** Our findings present a distinct molecular class of pediatric malignant gliomas largely overlapping with the recently reported bithalamic gliomas characterized by *EGFR* alteration, but additionally showing a broader spectrum of *EGFR* alterations and tumor localization. Global H3K27me3 loss in this group appears to be mediated by either H3 K27 mutation or *EZHIP* overexpression. EGFR inhibition may represent a potential therapeutic strategy in these highly aggressive gliomas.

**Key points:**

1. This study confirms a distinct new subset of pediatric diffuse midline glioma with H3K27me3 loss, with or without H3 K27 mutation
2. The poor outcome of these tumors is in line with the broader family of pediatric diffuse midline gliomas with H3 K27 mutation or *EZHIP* overexpression
3. Frequent *EGFR* alterations in these tumors may represent a therapeutic target in this subset

**Importance of the Study:** Malignant astrocytic gliomas in children show a remarkable biological and clinical diversity. Here, we highlight a distinct molecular class of pediatric malignant gliomas characterized by *EGFR* alteration and global H3K27me3 loss that appears to be mediated by either H3 K27 mutation or *EZHIP* overexpression. EGFR inhibition may represent a potential therapeutic strategy in these highly aggressive gliomas.

## Introduction

Malignant astrocytic gliomas of World Health Organization (WHO) grade 3 or 4 are among the most common malignant central nervous system (CNS) tumors in childhood^1^, and show remarkable biological and clinical heterogeneity. Despite common histopathological characteristics, pediatric malignant gliomas significantly differ from their adult counterparts in terms of molecular features, anatomical locations, and clinical outcomes^2–5^. Over the past decade, (epi-)genomic analyses have delivered broad insights into the molecular diversity of pediatric gliomas and enabled a subdivision into several biologically and clinically distinct entities defined by typical genetic alterations and DNA methylation profiles such as K27M or G34R/V mutations in histone 3, amongst others^6–8^.

In a recent study by Mondal et al.^9^, small in-frame insertions or missense mutations in the *EGFR* gene were observed in a subset of pediatric bithalamic diffuse gliomas, that also appeared to share a unique DNA methylation pattern and global loss of H3 K27me3. In order to further corroborate these findings, we used a screening approach based on unsupervised analysis of genome-wide DNA methylation profiling data of these initially reported cases alongside a large set of other CNS tumors (n > 30,000). This analysis revealed a molecularly distinct group of 58 tumors (including five of the cases initially described by Mondal et al.), for which further molecular workup was performed using targeted next-generation DNA sequencing.

## Materials and methods

### Sample collection

Tumor samples and retrospective clinical data were provided by multiple international collaborating centers and collected at the Department of Neuropathology of the University Hospital Heidelberg (Germany). Case selection was based on genome-wide DNA methylation screening using DNA methylation data of the cases initially described by Mondal et al.^9^ as a reference, which revealed a molecularly distinct group of tumors comprising five of those tumors with an additional 53 cases. Additionally, DNA methylation data of numerous well-characterized reference samples representing central nervous system (CNS) tumors of known histological and/or molecular subtype were used for comparative analyses^10,11^. Detailed descriptions of the reference DNA methylation classes are outlined under (https://www.molecularneuropathology.org). Analysis of tissue and clinical data was performed in accordance with local ethics regulations. Clinical details of the patients are listed in Supplementary Table 1.

### DNA methylation array processing and copy number profiling

Genomic DNA was extracted from fresh-frozen or formalin-fixed and paraffin-embedded (FFPE) tissue samples. DNA methylation profiling of all samples was performed using the Infinium MethylationEPIC (850k) BeadChip (Illumina, San Diego, CA, USA) or Infinium HumanMethylation450 (450k) BeadChip array (Illumina) as previously described^10^. Data of the samples of Mondal et al., were retrieved from Gene Expression Omnibus (GEO, https://www.ncbi.nlm.nih.gov/geo/) accession GSE140124. All computational analyses were performed in R version 3.3.1 (R Development Core Team, 2016; https://www.R-project.org). Copy-number variation analysis from 450k and EPIC methylation array data was performed using the conumee Bioconductor package version 1.12.0. Raw signal intensities were obtained from IDAT-files using the minfi Bioconductor package version 1.21.4^12^. Illumina EPIC samples and 450k samples were merged to a combined data set by selecting the intersection of probes present on both arrays (combineArrays function, minfi). Each sample was individually normalized by performing a background correction (shifting of the 5% percentile of negative control probe intensities to 0) and a dye-bias correction (scaling of the mean of normalization control probe intensities to 10,000) for both color channels. Subsequently, a correction for the type of material tissue (FFPE/frozen) and array type (450k/EPIC) was performed by fitting univariable, linear models to the log2-transformed intensity values (removeBatchEffect function, limma package version 3.30.11). The methylated and unmethylated signals were corrected individually. Beta-values were calculated from the retransformed intensities using an offset of 100 (as recommended by Illumina). All samples were checked for duplicates by pairwise correlation of the genotyping probes on the 450k/850k array. To perform unsupervised non-linear dimension reduction, the remaining probes after standard filtering^10^ were used to calculate the 1-variance weighted Pearson correlation between samples. The resulting distance matrix was used as input for t-SNE analysis (t-distributed stochastic neighbor embedding; Rtsne package version 0.13). The following non-default parameters were applied: theta□=□0, □=□F, max_iter□=□10,000 perplexity□=□15.

### Targeted next⍰generation DNA sequencing and mutational analysis

Capture-based next-generation DNA sequencing for cases with sufficient material (n=18) was performed on a NextSeq 500 instrument (Illumina) as previously described^13^ using a custom brain tumor panel covering the entire coding and selected intronic and promoter regions of 130 or 160 genes of particular relevance in central nervous system tumors. Reads were aligned against the reference genome (GRch37). Additionally, for 12 cases sequencing data generated locally at the initial handling institute were integrated in the present study.

### Gene expression profiling

Tumor samples with sufficient high-quality RNA (n=5) were analyzed on the Affymetrix GeneChip Human Genome U133 Plus (v.2.0) Array (Affymetrix, Santa Clara, USA) at the Microarray Department of the University of Amsterdam, the Netherlands or the Genomics and Proteomics Core Facility of the German Cancer Research Center (DKFZ), as previously described^14^. Expression data were normalized using the MAS5.0 algorithm. Expression levels of *EZHIP* were compared to published reference cases of H3 K27M-mutant and wildtype diffuse midline glioma^14,15^.

### Immunohistochemistry

Immunohistochemistry was performed on a Ventana BenchMark ULTRA Immunostainer using the ultraView Universal DAB Detection Kit (Ventana Medical Systems, Tucson, AZ, USA). Histone H3 lysine 27 trimethylated protein was detected using an anti-H3K27me3 antibody (rabbit polyclonal, Millipore, Burlington, MA, USA) diluted 1:500.

### Statistical analysis

Data on overall survival could be retrospectively retrieved for 14 patients. Overall survival was defined as time from initial diagnosis to patient death or last follow-up. Survival data were analyzed by Kaplan–Meier analysis and compared by log rank test using GraphPad Prism 8 (GraphPad Software, La Jolla, CA, USA). p values < 0.05 were considered significant.

## Results

### DNA methylation profiling reveals an epigenetically distinct group of pediatric malignant gliomas

We first investigated the distribution of the samples from the study of Mondal et al. based on unsupervised analysis of genome-wide DNA methylation profiling data alongside a large set of CNS tumors. Notably, this revealed a group of 53 tumor samples, clearly enriched for tumors of pediatric patients (see below), which formed a highly distinct DNA methylation class together with five of the recently published bithalamic glioma cases. A more focused t-distributed stochastic neighbor embedding (t-SNE) analysis of DNA methylation patterns of these 58 samples together with 516 well-characterized CNS neoplasms from the reference series encompassing various other types of pediatric and adult malignant astrocytic gliomas consistently confirmed the distinct nature of this novel group (Fig.1). The remaining five cases from the cohort described by Mondal et al. showed a DNA methylation profile matching that of the pedRTK2 subgroup, which has also been shown to have a high frequency of *EGFR* alterations^11^.

**Fig. 1.**
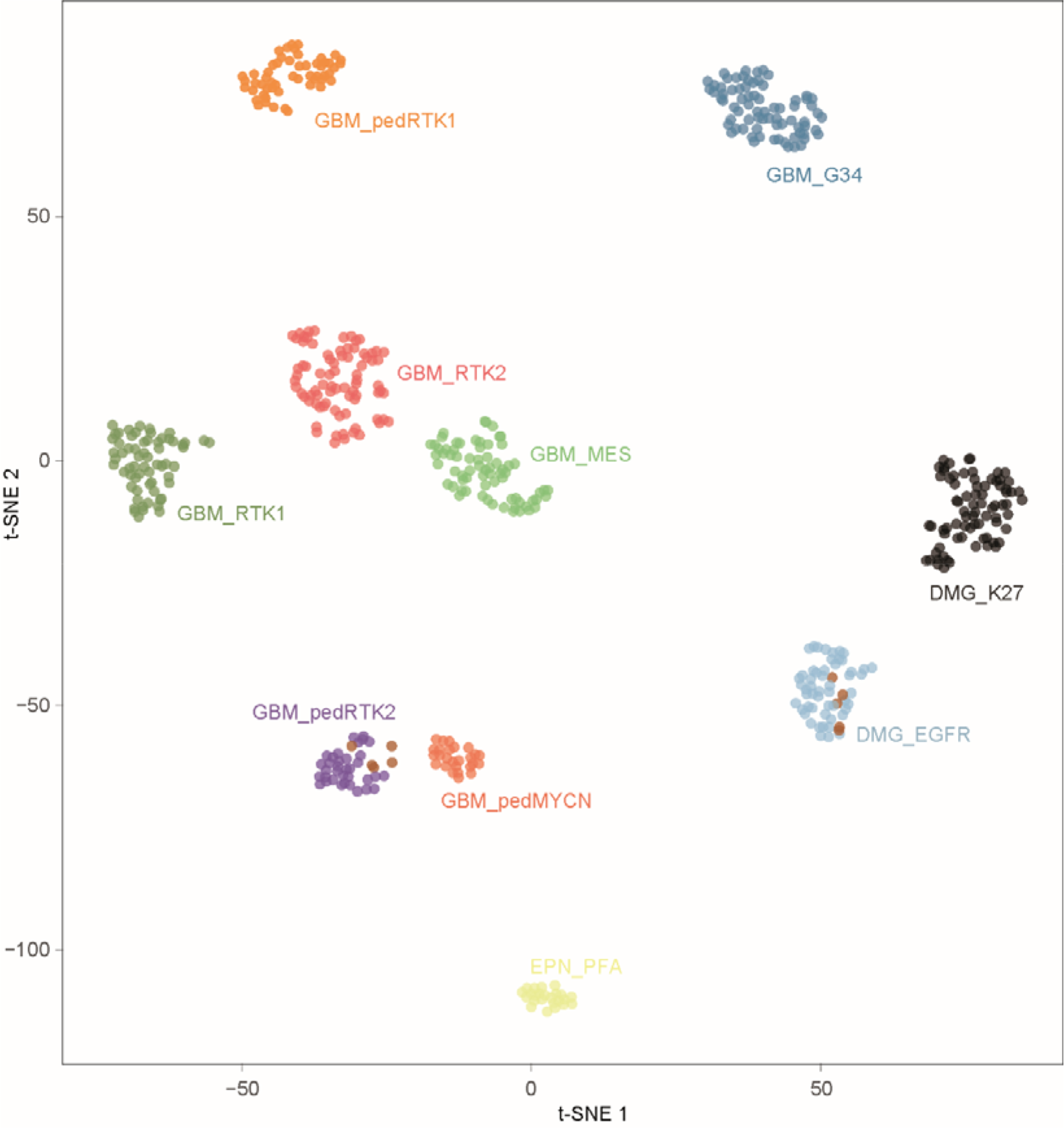
t-distributed stochastic neighbour embedding (t-SNE) analysis of DNA methylation profiles of the 58 gliomas investigated (DMG_EGFR; ten of the cases included from the Mondal et al. cohort (UCSF) are highlighted in brown) alongside selected reference samples. Reference DNA methylation classes: glioblastoma IDH-wildtype, subclass RTK 1 (GBM_RTK1); glioblastoma IDH-wildtype, subclass RTK 2 (GBM_RTK2); glioblastoma IDH-wildtype, subclass mesenchymal (GBM_MES); glioblastoma IDH-wildtype, pediatric RTK 1 (GBM_pedRTK1); glioblastoma IDH-wildtype, pediatric RTK 2 (GBM_pedRTK2); glioblastoma IDH-wildtype, pediatric MYCN (GBM_pedMYCN); diffuse midline glioma H3 K27M-mutant (DMG_K27M); glioblastoma IDH-wildtype, H3.3 G34-mutant (GBM_G34); ependymoma, posterior fossa group A (EPN_PFA).

### *EGFR* alteration and H3K27me3 loss are frequent features in this group of pediatric malignant gliomas

Analysis of copy number profiles (CNPs) derived from DNA methylation data revealed focal *EGFR* gene amplifications in 16 of the 58 tumor samples within the cohort (27%; Supplementary Fig. 1a) as well as several additional chromosomal aberrations including chromosome 7 gain, chromosome 1q gain, and chromosome 6q loss, each present in approximately half of all cases (Supplementary Fig. 1b). In light of the high frequency of *EGFR* amplifications and previously reported mutations/indels in the bithalamic cases, we next performed DNA sequencing of tumor samples with sufficient material (n=18). For an additional 12 cases, sequencing data provided by the original case submitter were integrated. In 15/30 analyzed tumors, a missense mutation in *EGFR* was detected, including nine tumors with a hotspot p.A289V/T substitution and single cases showing p.L62R/p.V774M, p. L861Q/p.T263P, p.Q276L, p.P719F, p.G598V, and p.T785P substitutions, respectively. Five further tumors showed an in-frame insertion within exon 20 of *EGFR,* including three of the tumors initially described by Mondal et al.^*9*^. Five of the *EGFR*-mutant tumors harbored a concomitant amplification of *EGFR.* Additionally, 18 of 30 tumors (62%) harbored a mutation or deletion of the tumor suppressor gene *TP53.* Subsets of tumors harbored additional hotspot mutations in genes encoding members of the mitogen-activated protein kinase (MAPK) and/or phosphoinositide 3-kinase (PI3K) signaling pathways including four tumors with a *PIK3CA* mutation and four tumors with either a *BRAF* V600E or a *NF1* mutation. Details are given in Figure 2 and Supplementary Table 1.

**Fig. 2.**
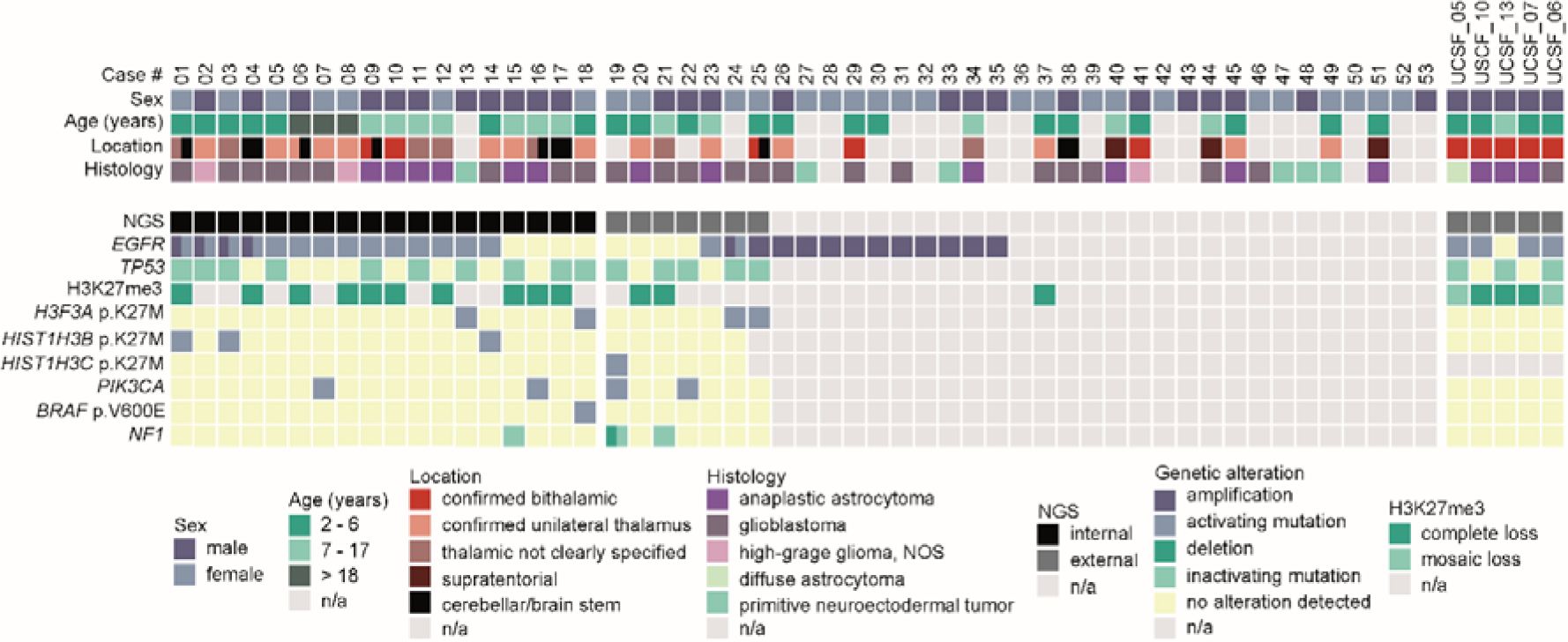
Clinicopathological characteristics and recurrent genetic alterations of the 58 gliomas.

Notably, eight of the 30 sequenced cases (27%) showed a K27M mutation in H3.1 or H3.3. Immunohistochemical staining for histone H3 lysine 27 trimethylation (H3K27me3) showed global loss of nuclear H3 K27me3 expression in tumor cells of all cases that could be evaluated (n=18; one of them showing an H3.3 K27M mutation; Fig. 3). Interestingly, analysis of *EZHIP* expression in five of the samples with available gene expression data indicated an overexpression in both of the H3-wildtype tumors compared to tumors with H3 K27M mutation (Fig. 4), in line with an alternative mechanism of PRC2 inhibition in glioma by *EZHIP* overexpression^15^. Despite these findings, no striking similarity was observed with the DNA methylation profile of other H3 K27M-mutant tumors including diffuse midline gliomas / diffuse intrinsic pontine gliomas as well as rare posterior fossa type A ependymomas (Fig. 1).

**Fig. 3.**
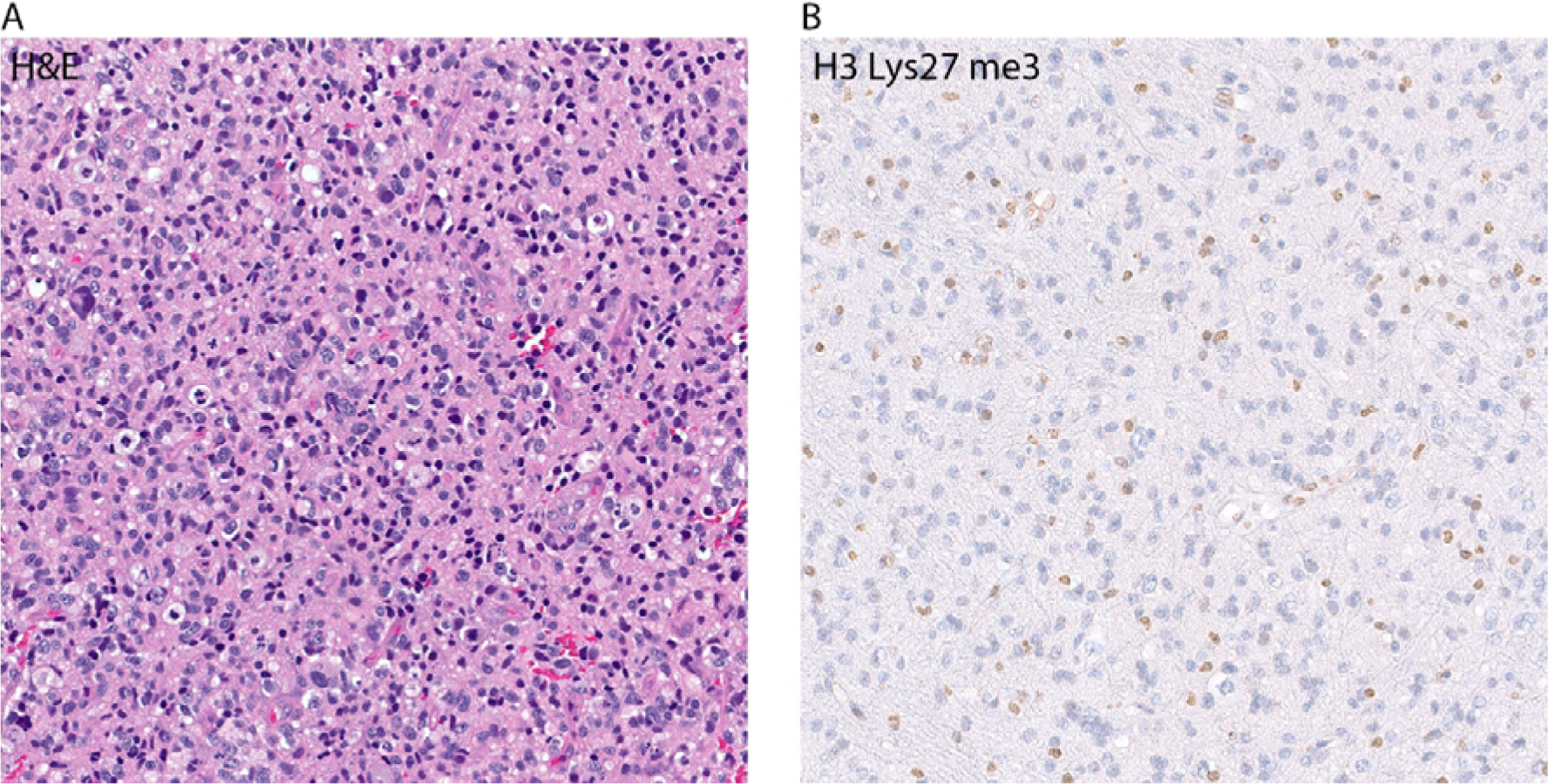
H&E staining of one of the gliomas included in the investigated cohort (case #1) revealing a pleomorphic astrocytic neoplasm with mitotic figures (A). Immunohistochemical staining for histone H3 lysine 27 trimethylation (H3K27me3) showing a loss of nuclear expression in the tumor cells (B).

**Fig. 4.**
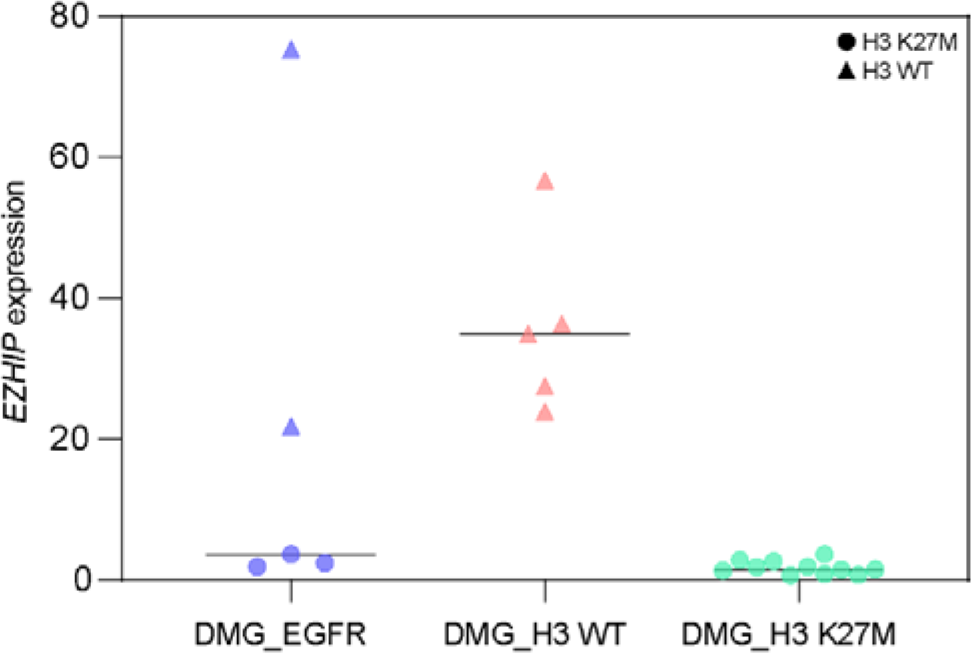
Distribution of normalized *EZHIP* gene expression level in the investigated glioma cohort in comparison to other diffuse midline gliomas showing that H3 K27M-wildtype tumors exhibit higher expression of *EZHIP.*

### Clinical characteristics and outcome

Limited clinical data were available for 36 of the patients. Tumors in most patients were histopathologically diagnosed as malignant astrocytic gliomas corresponding to glioblastoma or anaplastic astrocytoma (Fig. 2). Median patient age at the time of diagnosis was 8.2 years (range: 1–43 years) and the sex distribution was equal (male:female ratio 1:1.1). The majority of the tumors (n=31) were located in the thalamic region (Fig. 5), with five of the new tumors showing a clear bithalamic growth pattern (in addition to the five tumors reported by Mondal et al.). A small subset of tumors arose in the cerebral hemisphere (n=3), brain stem (n=2), and cerebellum (n=1). Retrospective outcome data concerning overall survival (OS) were available for 14 patients. Analysis of OS of these patients in comparison to children with diffuse midline glioma, H3 K27M-mutant (n=46) revealed a similarly poor outcome, and would support a putative assignment of WHO grade 4 to these tumors (Fig. 6).

**Fig. 5.**
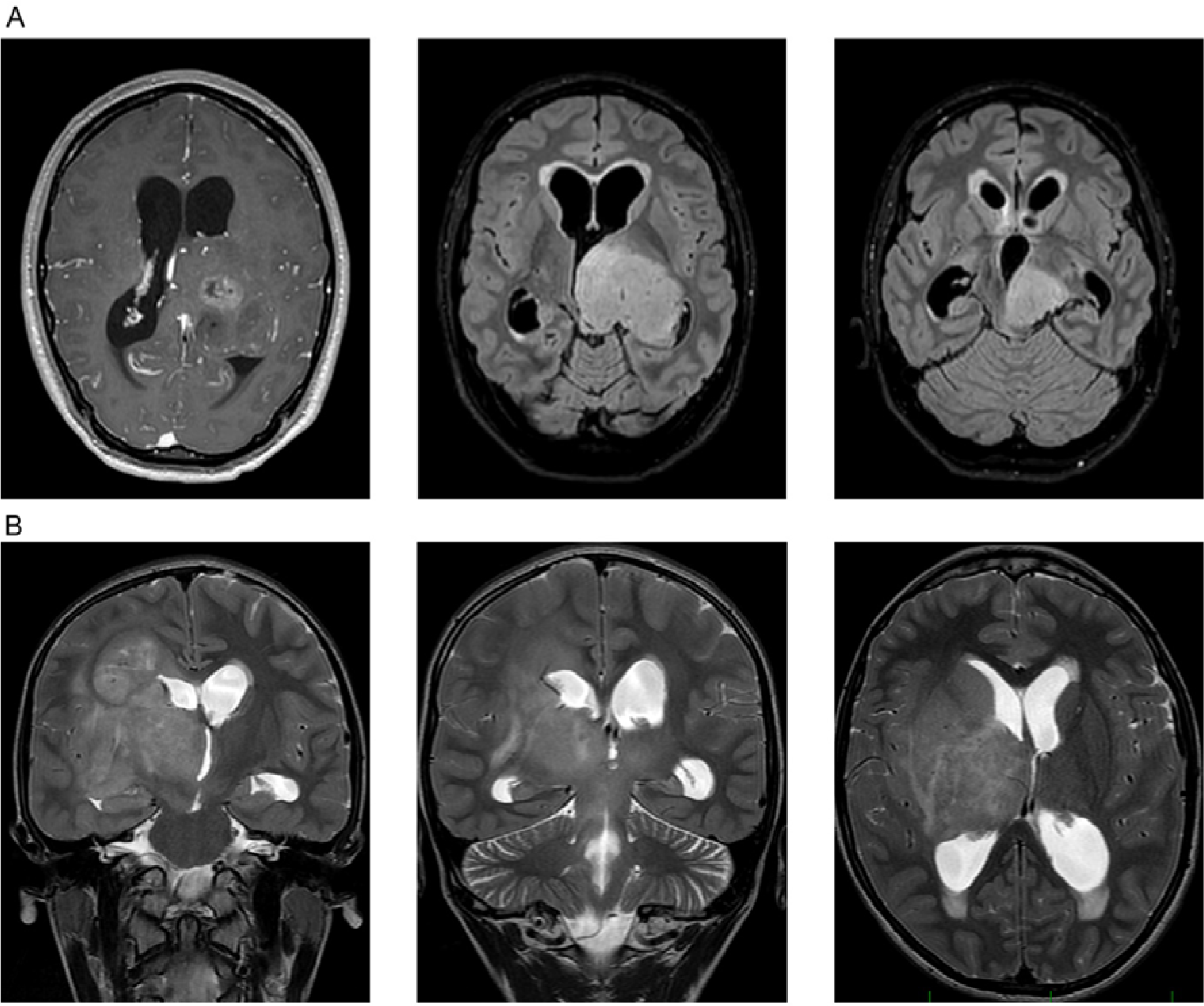
MR images from two representative patients of the investigated cohort showing a 21-years-old male patient (case #6) with a mass in the left thalamus, extending into mesencephalon and left cerebellar peduncle (A) and a 12-years-old boy (case #21) with a mass in the right thalamus with diffuse infiltration of the right hemisphere and T2 changes also in the contralateral hemisphere (B).

**Fig. 6.**
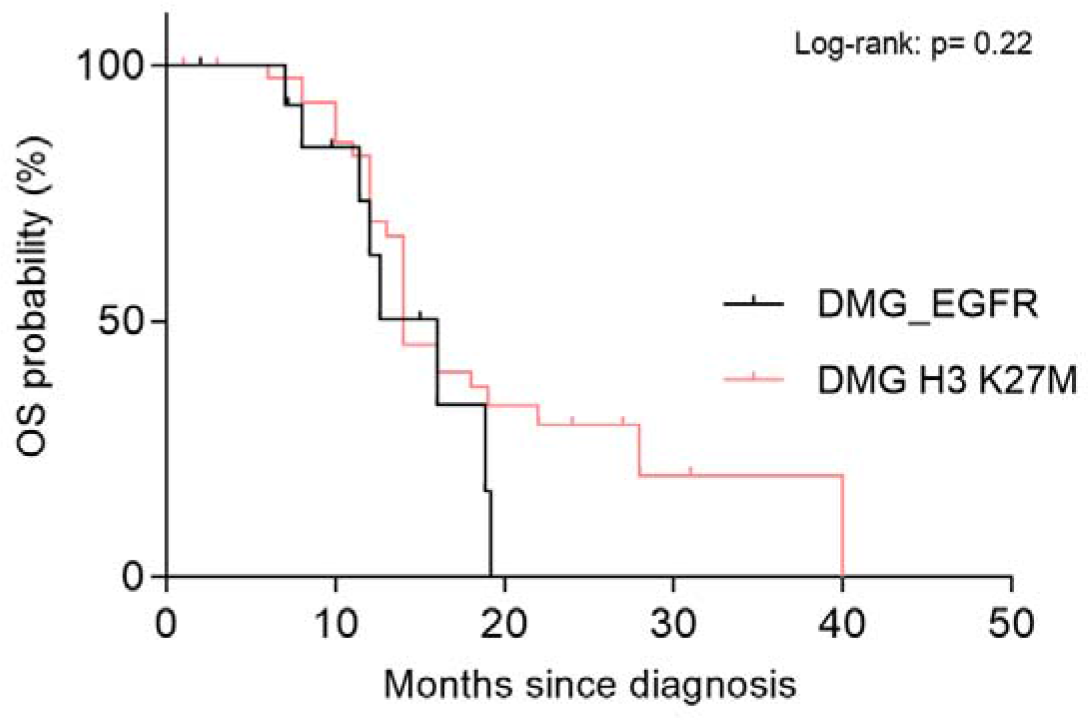
Kaplan–Meier curves for overall survival (OS) of 14 patients from the investigated cohort (DMG_EGFR) for whom OS data were available. OS of 14 patients is compared to OS of 46 patients with diffuse midline glioma, H3 K27M-mutant, WHO grade 4 (DMG H3 K27M) diagnosed at the University Hospital Heidelberg.

## Discussion

Here, we provide further evidence for the remarkable biological diversity of malignant astrocytic gliomas in children by confirming the existence of an epigenetically distinct subset of diffuse gliomas with predominantly thalamic location that shows frequent gene amplification and/or mutation of the *EGFR* gene and loss of H3K27me3. Overall, this group shows significant similarities with the class of “pediatric bithalamic gliomas” that has recently been reported by Mondal et al.^9^. Although most of the tumors in our series were located in the thalamic region, a small subset occurred in other, mostly midline CNS locations.

In addition, we expand on the spectrum of *EGFR* alterations observed in this group, with a high number of cases showing *EGFR* amplification and/or other missense mutations in *EGFR* besides the previously described variants^9^. Interestingly, only two additional tumors in our series (further to the 3/5 that were included from the Mondal et al.^9^ cohort) harbored the small in-frame insertions/duplications within exon 20 of *EGFR* which was a high-frequency event in the initial bithalamic glioma cohort. However, it should also be noted that sequencing data were available for only half of the novel cases in our larger cohort. Alterations of *EGFR* have been reported rarely in other types of pediatric malignant gliomas, but seem to be enriched particularly within the DNA methylation class glioblastoma IDH-wild-type, RTK III in the current version of the Heidelberg DNA methylation classifier (broadly equivalent to ‘pedRTK2’ according to Korshunov et al., 2017)^11^. The frequency of *EGFR* alterations in the present subset of pediatric malignant gliomas, however, is much higher than that reported for any other type of glioma except the RTK II subgroup of IDH-wildtype glioblastoma in adults, which also carries frequent *EGFR* alterations, in particular *EGFR* amplification^8^. These findings further identify a subset of pediatric malignant glioma with a potential therapeutic target, since potent EGFR inhibitors are available and previous work in a patient-derived orthotopic xenograft model has shown that pediatric gliomas with *EGFR* amplifications may respond to targeted inhibitors^16^. However, this hypothesis needs clinical validation as several clinical trials of EGFR-targeting approaches have previously failed in clinical trials for adult glioblastoma^17–19^.

In line with the previous report^9^, a high percentage of tumors in the present cohort harbored additional *TP53* mutations and a smaller subset showed H3 K27M mutations. These observations interestingly reveal that not all diffuse gliomas involving midline structures of the CNS with histone H3 K27M mutation align by DNA methylation profile with the reference DNA methylation class “diffuse midline glioma, H3 K27M-mutant” (which does not typically feature *EGFR* alterations)^3^. This new group of diffuse midline gliomas with uniform H3K27me3 loss and frequent *EGFR* alteration can also harbor H3 K27M mutation in a subset. Distinction between these two tumor types cannot be reliably determined by histologic features or H3 K27M mutation status alone, and thus requires assessment of *EGFR* copy number and mutation status or DNA methylation signature at present. Notably, not all H3 K27M mutations were comparably covered by the different panel sequencing approaches integrated in the present study so that the frequency of alterations might be underestimated.

A major limitation of our study is the relatively low number of tumor samples with available tissue for a more comprehensive molecular work-up. Further analyses are needed to reveal the full spectrum of genetic alterations within this group of tumors. Considering the features identified in the recent previous study^9^ and evaluating the overlaps and additional findings in the current set, we propose the broader designation of ‘diffuse midline glioma, H3K27me3-lost and *EGFR*-altered’ which would include the majority of bithalamic diffuse gliomas as well as a subset of unilateral thalamic gliomas.

Intriguingly, loss of H3K27me3 can now be regarded as unifying feature of three molecular classes of diffuse astrocytic glioma with predominant midline location: The “typical” diffuse midline glioma with H3K27M mutation, diffuse midline glioma with EZHIP overexpression, and the series presented here. Thus, integrating these three to a novel overarching tumor type “diffuse midline glioma, H3K27me3 loss” and considering the three molecular classes as subtypes could be a reasonable approach to implement these findings into the WHO classification.

In summary, we provide further evidence for a subset of pediatric-type malignant glioma that is preferentially located in the thalamic region and shows frequent alteration of *EGFR,* which might provide future options for targeted therapeutic approaches. Also, the occurrence of H3 K27M mutations in this glioma subgroup has implications for classification. Assessment of H3 mutation status alone is not sufficient to distinguish ‘typical’ diffuse midline glioma, H3 K27M-mutant tumors (which occur much more frequently based on the larger methylation database) from this novel glioma subtype with loss of nuclear H3K27me3 expression and frequent *EGFR* alteration. Although this diagnostic distinction has limited implications for patient outcome at present, our results suggest that a more comprehensive molecular workup of diffuse thalamic gliomas may eventually become clinically relevant, e.g. if shown to be associated with distinct responses to molecularly targeted therapeutic approaches.

## Acknowledgements

We thank L. Dörner, L. Hofmann, and M. Schalles for skillful technical assistance and the microarray unit of the DKFZ Genomics and Proteomics Core Facility for providing Illumina DNA methylation array-related services.

## References

1. Ostrom QT, Cioffi G, Gittleman H, et al. CBTRUS Statistical Report: Primary Brain and Other Central Nervous System Tumors Diagnosed in the United States in 2012-2016. Neuro Oncol. 2019; 21(Suppl 5):v1–v100.

2. Jones C, Baker SJ. Unique genetic and epigenetic mechanisms driving paediatric diffuse highgrade glioma. Nat Rev Cancer. 2014; 14(10).

3. Mackay A, Burford A, Carvalho D, et al. Integrated Molecular Meta-Analysis of 1,000 Pediatric High-Grade and Diffuse Intrinsic Pontine Glioma. Cancer Cell. 2017; 32(4):520–537 e525.

4. Paugh BS, Qu C, Jones C, et al. Integrated molecular genetic profiling of pediatric high-grade gliomas reveals key differences with the adult disease. J Clin Oncol. 2010; 28(18):3061–3068.

5. Sturm D, Bender S, Jones DT, et al. Paediatric and adult glioblastoma: multiform (epi)genomic culprits emerge. Nat Rev Cancer. 2014; 14(2):92–107.

6. Schwartzentruber J, Korshunov A, Liu XY, et al. Driver mutations in histone H3.3 and chromatin remodelling genes in paediatric glioblastoma. Nature. 2012; 482(7384):226–231.

7. Wu G, Broniscer A, McEachron TA, et al. Somatic histone H3 alterations in pediatric diffuse intrinsic pontine gliomas and non-brainstem glioblastomas. Nat Genet. 2012; 44(3):251–253.

8. Sturm D, Witt H, Hovestadt V, et al. Hotspot mutations in H3F3A and IDH1 define distinct epigenetic and biological subgroups of glioblastoma. Cancer Cell. 2012; 22(4):425–437.

9. Mondal G, Lee JC, Ravindranathan A, et al. Pediatric bithalamic gliomas have a distinct epigenetic signature and frequent EGFR exon 20 insertions resulting in potential sensitivity to targeted kinase inhibition. Acta Neuropathol. 2020; 139(6):1071–1088.

10. Capper D, Jones DTW, Sill M, et al. DNA methylation-based classification of central nervous system tumours. Nature. 2018; 555(7697):469–474.

11. Korshunov A, Schrimpf D, Ryzhova M, et al. H3-/IDH-wild type pediatric glioblastoma is comprised of molecularly and prognostically distinct subtypes with associated oncogenic drivers. Acta Neuropathol. 2017; 134(3):507–516.

12. Aryee MJ, Jaffe AE, Corrada-Bravo H, et al. Minfi: a flexible and comprehensive Bioconductor package for the analysis of Infinium DNA methylation microarrays. Bioinformatics. 2014; 30(10):1363–1369.

13. Sahm F, Schrimpf D, Jones DT, et al. Next-generation sequencing in routine brain tumor diagnostics enables an integrated diagnosis and identifies actionable targets. Acta Neuropathol. 2016; 131(6):903–910.

14. International Cancer Genome Consortium PedBrain Tumor P. Recurrent MET fusion genes represent a drug target in pediatric glioblastoma. Nat Med. 2016; 22(11):1314–1320.

15. Castel D, Kergrohen T, Tauziede-Espariat A, et al. Histone H3 wild-type DIPG/DMG overexpressing EZHIP extend the spectrum diffuse midline gliomas with PRC2 inhibition beyond H3-K27M mutation. Acta Neuropathol. 2020; 139(6):1109–1113.

16. Brabetz S, Leary SES, Grobner SN, et al. A biobank of patient-derived pediatric brain tumor models. Nat Med. 2018; 24(11):1752–1761.

17. Lassman AB, Rossi MR, Raizer JJ, et al. Molecular study of malignant gliomas treated with epidermal growth factor receptor inhibitors: tissue analysis from North American Brain Tumor Consortium Trials 01-03 and 00-01. Clin Cancer Res. 2005; 11(21):7841–7850.

18. Hegi ME, Diserens AC, Bady P, et al. Pathway analysis of glioblastoma tissue after preoperative treatment with the EGFR tyrosine kinase inhibitor gefitinib--a phase II trial. Mol Cancer Ther. 2011; 10(6):1102–1112.

19. Weller M, Butowski N, Tran DD, et al. Rindopepimut with temozolomide for patients with newly diagnosed, EGFRvIII-expressing glioblastoma (ACT IV): a randomised, double-blind, international phase 3 trial. Lancet Oncol. 2017; 18(10):1373–1385.

